# MOGAT: An Improved Multi-Omics Integration Framework Using Graph Attention Networks

**DOI:** 10.1101/2023.04.01.535195

**Authors:** Raihanul Bari Tanvir, Md Mezbahul Islam, Masrur Sobhan, Dongsheng Luo, Ananda Mohan Mondal

## Abstract

Integration of multi-omics data holds great promise for understanding the complex biology of diseases, particularly Alzheimer’s, Parkinson’s, and cancer. However, the integration is challenging due to the high dimensionality and complexity of the data. Traditional machine learning methods are not well-suited for handling the complex relationships between different types of omics data. Many models were proposed that utilize graph-based learning models to extract hidden representations and network structures from different omics data to enhance cancer prediction, patient categorization, etc. The existing graph neural network-based (GNN-based) multi-omics approaches for cancer subtype prediction have three shortcomings: (a) Do not consider all types of omics data, (b) Fail to determine the relative significance of the neighboring nodes (in this case, samples or patients) when it comes to downstream analyses, such as subtype classification, patient stratification, etc., and (c) Use the same approach for generating initial graphs for different omics data. To overcome these shortcomings, we present MOGAT, a novel multi-omics integration approach, leveraging a graph attention network (GAT) model that incorporates graph-based learning with an attention mechanism. MOGAT utilizes a multi-head attention mechanism that can more efficiently extract information for a specific sample by assigning unique attention coefficients to its neighboring samples. To evaluate the performance of MOGAT, we explored its capability via a case study of predicting breast cancer subtypes. Our results showed that MOGAT performs better than the state-of-the-art multi-omics integration approaches.

## 1 Introduction

The advent of next-generation sequencing NGS technologies has led to the generation of a vast amount of multi-omics data, posing major challenges in data analysis and integration. Integrating multi-omics data is crucial for gaining a comprehensive understanding of complex disease like Alzheimer’s [1], Parkinson’s [2], [3] and cancer [4]. However, it is a difficult task that requires advanced computational methods. There is a need for the development of new analytical tools and methods that can effectively extract biologically relevant information from multi-omics data and integrate it into a comprehensive understanding of the disease. Despite the challenges of high dimensionality and complexity of data, the integration of multi-omics data holds great potential for understanding the biology of cancer.

Many models were proposed that utilize graph-based learning models to extract hidden representations and graph structures from different omics data to enhance the understanding of Alzheimer’s, Parkinson’s, and cancer prediction, patient categorization, etc. Wang et al. [1] used multi-omics integration for Alzheimer’s disease patient classification. Researchers also used multi-omics integration to find molecular biomarkers [3] and disease classification [2] for Parkinsons’s disease. A lot of multi-omics research have been conducted in predicting cancer subtypes and patient categorization. Li et al. [5] utilize graph convolutional network (GCN) [6] to classify 28 different cancer types from pan-cancer data using gene expression and copy number alteration as features and three knowledge networks as the input graphs, including gene-gene interaction (GGI) networks, protein-protein interaction (PPI) networks, and gene co-expression networks. Zhou et al. [7] use gene expression, DNA methylation, and miRNA expression as features for multi-omics analysis. They use anchors to derive sample similarity networks and graph convolutional autoencoder for clustering cancer samples to identify novel subtypes for breast, brain, colon, and kidney cancer. Guo et al. [8] use GCN by taking the PPI network as a graph and gene expression, copy number alterations, and DNA methylation as features. Finally, they applied attention on top of embeddings generated by GCN to classify breast cancer subtypes. Li et al. proposed MoGCN [9], which uses an autoencoder for feature extraction and similarity network fusion to construct the patient similarity network. It applies GCN to classify breast cancer subtypes and pan-kidney cancer type classification using gene expression, copy number alterations, and phase protein array data as input. M-GCN [10] is another multi-omics framework based on GCN to classify cancer subtypes of breast and stomach cancer. They use the Hilbert– Schmidt independence criterion-based least absolute shrinkage and selection operator (HSIC LASSO) to select the molecular subtype-related transcriptomic features and then use those features to construct a patient similarity graph applying Pearson’s correlation. For multi-omics data, it takes gene expression, single nucleotide variation, and copy number alterations as input. MOGONET [1] takes gene expression, DNA methylation, and miRNA as input. It uses, unlike other methods, GCN to learn omics-specific embeddings and uses network and node features for a particular omics data. Then it combines the embeddings using a view correlation discovery network (VCDN) to classify cancer subtypes for breast, brain, and pan-kidney cancer. The SUPREME[11] method utilizes GCN for analyzing breast cancer subtypes. It integrates seven types of data - gene expression, miRNA expression, DNA methylation, single nucleotide variation, copy number alteration, co-expression module eigengenes, and clinical data - for constructing the network and determining node features. Additionally, it combines GCN embeddings with node features and employs a Multi-layer Perceptron (MLP) as a classifier.

In summary, the existing GNN-based muti-omics integration approaches to predict cancer subtypes apply GCN to extract the salient features from different omics data. However, GCN-based frameworks cannot determine the relative significance of neighboring samples regarding downstream analysis, including cancer subtype prediction, patient stratification, etc. It is also noticeable that none of the existing studies considered long non-coding RNA (lncRNA) expression data in multi-omics integration. But lncRNAs play important regulatory roles in various cellular processes, including gene expression and epigenetic regulation [12]–[16]. This research presents MOGAT, a novel multi-omics integration-based cancer subtype prediction leveraging a graph attention network (GAT) [17] model that incorporates graph-based learning with an attention mechanism for analyzing multi-omics data. The proposed MOGAT utilizes a multi-head attention mechanism that can more efficiently extract information for a specific patient by assigning unique attention coefficients to its neighboring patients, i.e., getting the relative influence of neighboring patients in the patient similarity graph. We also included lncRNA expression in the multi-omics integration process, altogether eight different data types, including gene expression, miRNA expression, lncRNA expression, DNA methylation, single nucleotide variation, copy number alteration, co-expression module eigengenes, and clinical data. Based on our knowledge, our approach is the first approach to use GAT in multi-omics integration and used the most different types of omics data.

## 2 Materials and Methods

Figure 1 illustrates the diagram of our proposed multi-omics integration approach, MOGAT and its application in the prediction of breast cancer subtypes.

**Fig. 1.**
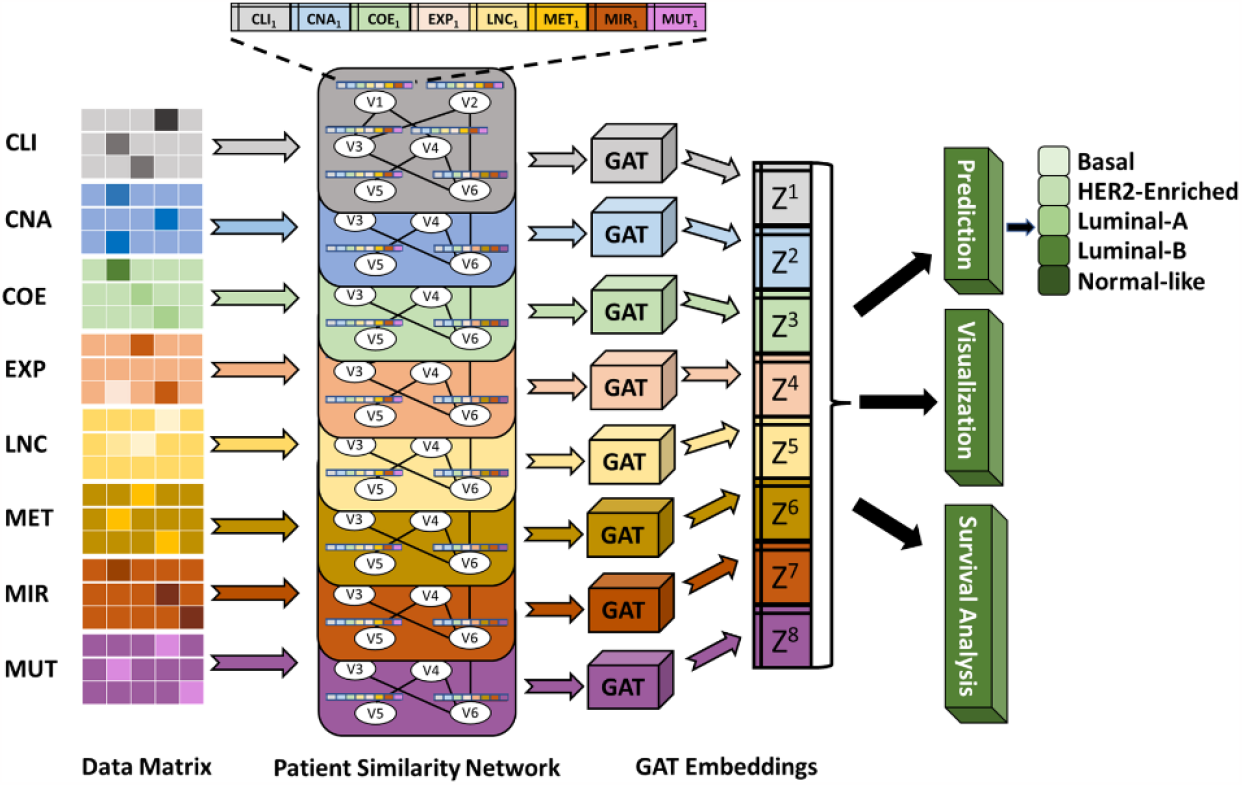
Illustration of MOGAT framework. The MOGAT framework processes patient similarity networks constructed from eight datatypes, including CLI (clinical), CNA (copy number alteration), COE (co-expression), EXP (mRNA expression), LNC (lncRNA expression), MET (DNA methylation), MIR (miRNA expression), and MUT (simple nucleotide variation). Nodes in each patient similarity network are annotated with features from eight data types. By applying GAT to each patient similarity network, the framework generates embeddings. These embeddings are then used for subtype prediction, visualization, and survival analysis.

### 2.1 Dataset Preparation and Preprocessing

To develop and investigate the MOGAT approach, we downloaded omics and clinical data for Breast Invasive Carcinoma (BRCA) from the GDC portal (https://portal.gdc.cancer.gov) of The Cancer Genome Atlas (TCGA). The RNAseq gene (mRNA, miRNA, and lncRNA) expression, DNA methylation, copy number variation, simple nucleotide variation, and clinical data were collected for this cohort. Table 1 shows the summary of the processed omics data. The preparation and preprocessing of different types of data are outlined below.

**Table 1.** The summary of each datatype. Row 1 (Original Features): number of original features, Row 2 (Cleaned Features): number of features after cleaning by filtering, Row 3 (Selected Features): number of features after applying feature selection approach, Boruta, Row 4 (All Tumor Samples): Number of tumor samples including duplicates, Row 5 (Unique Tumor Samples): number of tumor samples after removing duplicates, Row 6 (Common Samples): subtype distribution of the tumor samples common across all datatypes, and Row 7 (Network (Nodes, Edges)): number of nodes and edges for the patient similarity network for each datatype.

| Datatype | CLI | CNA | COE | EXP | LNC | MET | MIR | MUT |
| --- | --- | --- | --- | --- | --- | --- | --- | --- |
| Original Features | 31 | 28,918 | 40 | 19,962 | 16,901 | 25,978 | 1881 | 16,662 |
| Cleaned Features | - | - | - | 5,343 | 3,398 | - | 393 | - |
| Selected Features | 31 | 500 | 40 | 1000 | 500 | 1,000 | 306 | 200 |
| All Tumor Samples | 1,089 | 1,106 | 1,212 | 1,212 | 1,212 | 1,107 | 1,069 | 992 |
| Unique Tumor Samples | 1,089 | 1,096 | 1,076 | 1,076 | 1,076 | 1,097 | 1,057 | 969 |
| Common Samples | **920** Samples; Basal: 158 (17.24%) HER2: 73 (8.02%) LumA: 467 (50.65%) LumB: 188 (20.39%) NL: 34 (3.68%) |  |  |  |  |  |  |  |
| Network (Nodes, Edges) | (920, 2,398) | (920, 2,346) | (920, 2,122) | (920, 2,391) | (920, 2,218) | (920, 2,675) | (920, 2,108) | (920, 2,752) |

Note: CLI: clinical, CNA: copy number alteration, COE: co-expression, EXP: gene expression, LNC: lncRNA expression, MET: DNA methylation, MIR: miRNA expression, MUT: simple nucleotide variation. LumA: Luminal A, LumB: Luminal B, NL: Normal-like.

#### Features Based on Clinical Data

The clinical data consists of variables age, race, tumor status, tumor stage, menopause status, estrogen receptor status, progesterone receptor status, and HER2 receptor status. Except for age, the rest of the variables were converted into a one-hot vector. Finally, the number of features remained at 31.

#### Features Based on Copy Number Alterations

Copy number alteration data came as Mutation Annotation Format (MAF) files, each for one sample. Each row of the MAF file corresponds to a genomic coordinate for which the copy number values were observed as the mean segment value. These files were combined for a single MAF file for all patients containing 1,268,167 rows. Then, CNTools [18] was used to obtain gene-centric values from the segmented copy number variation data, and the number of genes was 28,918.

#### Features Based on Gene Co-Expression

For generating gene co-expression network, WGCNA [19] was used. Soft thresholding was used to construct the co-expression network. The optimal soft-threshold power beta was selected by evaluating its effect on the scale-free topology model fit R^2^. From the beta values from 1 to 30, 4 was the lowest while maintaining the high R^2^ values (threshold 0.90). The adjacency matrix was first transformed into a Topological Overlap Matrix (TOM) based similarity matrix for module detection. Then it was converted into a TOM-based dissimilarity matrix by subtracting from unity (1). This matrix was used as a distance metric to perform the average linkage hierarchical clustering algorithm, which outputs a dendrogram. Then Dynamic Tree Cut [20] method was performed for branch cutting to generate network modules. The minimum size of the modules was set at 40 genes.

#### Features Based on mRNA Expression

RNAseq expression data contains expression of 60,660 genes (including mRNAs, miRNAs, and lncRNAs) from which expression values of 19,962 mRNAs were isolated. The expression values were in FPKM (fragments per kilobase of exon per million mapped fragments). The mRNAs are filtered out if its expression values do not meet the threshold of FPKM >= 1 in >=15% of samples (as used in SUPREME), which resulted in 13,503 mRNAs. These mRNAs were used to perform differential gene expression analysis using DESeq2 [21]. After using the criteria of adjusted P-value <= 0.01, the number of remaining mRNAs was 5,343.

#### Features Based on lncRNA Expression

Applying the similar preprocessing approach used for mRNA, we isolated 3,398 lncRNAs and corresponding expression values from the original dataset of 60,660 gene expressions.

#### Features Based on miRNA Expression

The miRNAseq expression data were in reads per million (RPM) unit. The miRNAs are filtered out if its expression values do not meet the threshold of RPM>=1 in>= 30% of samples, which resulted in 393 miRNAs. Then differential gene expression analysis using adjusted P-value <= 0.01 resulted in 306 miRNAs.

#### Features Based on DNA Methylation

Both HumanMethylation 27k (HM27) and HumanMethylation 450k (HM450) data were collected for DNA methylation. For the HM27 data, the number of samples was 343, and the number of probes was 27,578. For the HM450 data, the numbers of samples and probes were 895 and 485,577, respectively. After combining both datasets by keeping the same probes, the samples and probes were 1,238 and 25,978, respectively.

#### Features Based on Simple Nucleotide Variation

For simple nucleotide mutation data, there were 992 samples, and each sample contained a different set of genes for which one or more mutations were observed. The sample mutation data were converted into a vector of genes, where one signifies a mutation occurred, and 0 signifies no mutation. The size of this vector is the union of all the genes from all samples, which is 16,662.

### 2.2 Network and Feature Matrix Construction

The input for the graph attention network is a network and the feature matrix for the nodes in the given network. The training, in this case, would be in single mode, where each sample corresponds to a node in the network, and the whole network represents a data type (mutation, gene expression, etc.). In this case, the patient similarity network is constructed where the node represents the patient or sample, and the edge represents the similarity between two patients. Before constructing the networks and the feature matrix for the nodes in the network, it was observed that some of the patients have more than one sample in different omics datasets. For example, a patient with ID: TCGAA7-A0DB contains three samples in gene expression data, mutation data, and miRNA expression data. So, to make sure each patient has one corresponding sample, an extra preprocessing step was taken. For gene expression data, miRNA expression data, co-expression data, DNA methylation, and copy number alterations data-the multiple samples corresponding to single patients were replaced with one sample by taking the average value for each feature. And for mutation data, since this is a binary 0/1 data, the Boolean OR operation was done instead of average for the patients with more than one sample.

For network construction, the same set of patients needed to exist for each patient similarity network corresponding to a data type. However, different omics dataset contains different numbers of samples. It was observed that 920 tumor samples exist in all types of omics data, so these 920 common tumor samples are used to create the patient similarity network and the feature matrix. The similarity matrix was created for each dataset to construct the patient similarity network. This similarity was computed using Pearson’s correlation for mRNA expression, miRNA expression, co-expression features, copy number alterations, and DNA methylation. Jaccard similarity was used instead of Pearson’s correlation network for mutation data, as it is binary data. The Gower metric [22] was used to compute patient similarity using clinical data since it combines categorical and continuous features. Then from each similarity matrix, the top 3 highest similar samples for each sample were selected as edges to construct the similarity network.

The Boruta package [23], a feature selection method based on the Random Forest algorithm, was used for the feature selection process. For feature matrix construction, a feature selection step was conducted for copy number alterations, mRNA expression, lncRNA expression, DNA methylation, and mutation, - as these omics data are high-dimensional low sample-size data. This step was not used for co-expression, miRNA expression, and one-hot encoded clinical features as they did not have many features as other counterparts.

### 2.3 Graph Attention Network

The utilized GAT model is based on the idea of the self-attention mechanism, where embeddings are created from eight different types of data (Table 1) with the assumption that samples with similar characteristics (such as gene expression or DNA methylation) are likely to have similar disease outcomes and are therefore related to each other. However, not all related samples should be given equal importance. Some samples may have a greater impact on the prediction or clustering of a target sample, which cannot be accurately determined by similarity metrics. To address this, the GAT model assigns varying levels of attention to the related (or, neighboring) samples of a target sample, allowing it to capture the significance of each one.

For each data type, let *n* be the number of samples or patients and *m* be the number of features (e.g., genes, miRNAs etc.). The input feature matrix is given by *X =* [*x*_1_, *x*_2_, …, *x*_*n*_], where *x* ∈ ℝ^1×*m*^ represents a sample vector. Let *A* be the *n* × *n* adjacency matrix build based on pairwise correlation between samples. The adjacency matrix is binarized, as it will be used to mask the attention coefficients in later part of the model. While generating the embedding of sample *x*_*i*_, the attention given to it from its neighbor *x*_*j*_ can be calculated as:

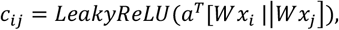

where W ∈ ℝ^*p*×*m*^ and *a* ∈ ℝ^2*p*×1^ are learnable weight parameters, which are shared across all samples and *p* is the embedding size; || symbol denotes the concatenation of two vectors; and LeakyReLU is the non-linear activation function. *c*_*ij*_ describes the importance of sample *j*’s feature to sample *i*. We then nomalize attention coefficients by applying a SoftMax function:

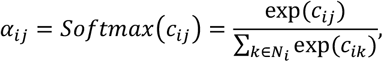

Where *N*_*i*_ is the set of neighboring nodes of sample *i*.With the normalized attention coefficients being the weights, a linear combination of input features is then used as the output representation for each data sample. Formally, we have:

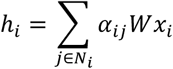

Where *h*_*i*_ is the output representation of sample *i*.

### 2.4 Classification

Embeddings generated after training the GATs were concatenated and used as input for classification. A multilayer perceptron (MLP) was used to classify breast cancer subtypes. The architecture, learning rate, and the number of epochs for the multilayer perceptron were selected based on a randomized grid search. The range of values is listed in Table 2.

**Table 2.** Hyperparameter Tuning for GAT, MLP, and LASSO. The type of the hyperparameter and their ranges of values used for tuning. Optimized hyperparameter values for GAT and LASSO are bolded. For MLP tuning, it has 255 set of optimized hyperparameter values, one for each combination of multi-omics data. Optimized parameters with eight data types are reported as bold.

| Hyperparameters | Values |
| --- | --- |
| GAT hidden layer dimensions | [128, 256, **512**, 1024] |
| GAT Learning Rate | [0.01, **0.001**, 0.0001] |
| GAT # of epochs | [100, **200**, 500] |
| MLP learning rate | [0.1, 0.01, **0.001**, 0.0001, 0.00001] |
| MLP hidden layer architecture | [(32), (64), (128), (256), (512), (32, 32),<br>(**64**, **32**), (128, 32), (256, 32)] |
| MLP # of epochs | [200, 500, **1000**, 1500] |
| LASSO regularizing factor $\alpha$ | [0.001,0.002,0.005,0.01,0.05,**1.0**] |

### 2.5 Training Process

For training the GATs, the architecture remained the same for all the datatypes, consisting of two graph attention layers. The hidden layer dimension for each GAT model and learning rate were selected based on grid search-based hyperparameter tuning. The ranges of values for hyperparameters are listed in Table 2.

The classification metrics, including accuracy, weighted-F1 score, and macro-F1 score were estimated to evaluate the performance of MOGAT model. The GATs were implemented using PyTorch Geometric 2.1.0 and MLP was implemented using PyTorch 1.13.0. The training and testing of deep learning models were performed on a GeForce GTX 1080 Ti 8 GPU, 256GB RAM, 28 core Intel Xeon CPU E5-2650.

### 2.6 Survival Analysis

Survival Analysis was performed using both raw features and GAT embeddings separately. Table 3 shows the number of raw features and GAT embeddings at different stages of survival analysis. The initial number of raw features and embeddings were 3,577 and 4,096, respectively. The LASSO regression was used to reduce the number of raw features and embeddings to 276 and 2,247, respectively, considering overall survival as output. The regularizing factor *α* for LASSO was selected using a Grid SearchCV approach. The list of values for *α* are given in Table 2. Next, a multivariate Cox Proportional Hazard (Cox-PH) regression analysis [24] was conducted using the selected features in the previous step. This technique examines the influence of multiple variables on the time it takes for a specific event to occur, in this case, death. In the Cox regression model, the coefficients of predictor variables (raw features or embeddings) are related to hazard, i.e., risk of death. A positive coefficient indicates a worse prognosis, and a negative coefficient indicates a protective effect of the variable with which it is associated. The hazard ratio associated with a predictor variable is given by the exponent of its coefficient, and the P-value shows the significance of association between the predictor variable (raw feature or embedding) and the risk of death, either increasing or decreasing. The significant predictor variables, 57 raw features and 542 embeddings, with a P-value < 0.05 were selected, and their Cox-coefficients were used to calculate the risk scores using raw features and embeddings, respectively.

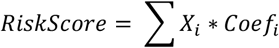

**Table 3.** The number of variables (Raw features and GAT embeddings) that remained at each step of the survival analysis process. Raw features are the sum of the reduced set of features from eight different data types (Table 1, Row: “Selected Features”). Each of the eight GAT embeddings reduced the raw features from 3,577 to 512.

| Item | Raw Features | Embeddings |
| --- | --- | --- |
| Initial Number of Variables | 3,577 | 8 * 512 = 4096 |
| After LASSO | 276 | 2247 |
| After Cox-PH P-value $\leq 0.05$ | 57 | 542 |

Where, *X*_*i*_ is the value of *i*-th predictor (raw feature or embedding) and *Coef*_*i*_ is the corresponding coefficient for the predictor obtained from the Cox-regression. This Risk Score is used to divide the cohort into low-risk and high-risk groups using median as the divider. Then Kaplan Meier [25] test and Logrank test [26] were performed and hazard ratio was calculated to see if the two groups are significantly distinguishable.

## 3 Result

### 3.1 Comparison of Performance

To assess the performance of the proposed MOGAT framework, we compared it with two state-of-the-art frameworks that integrate multi-omics data for subtype prediction, namely, MOGONET and SUPREME. Three omics data were used to show the performance comparison of MOGAT with MOGONET and SUPREME: gene expression, DNA methylation, and miRNA expression, as it was originally used in MOGONET. For the same three omics data, the test macro-F1 scores on seven (2^3^-1) combinations of three omics data were calculated and plotted in Figure 2(a). We observed that MOGAT has higher macro-F1 scores compared to MOGONET and SUPREME on the same test dataset.

**Fig. 2.**
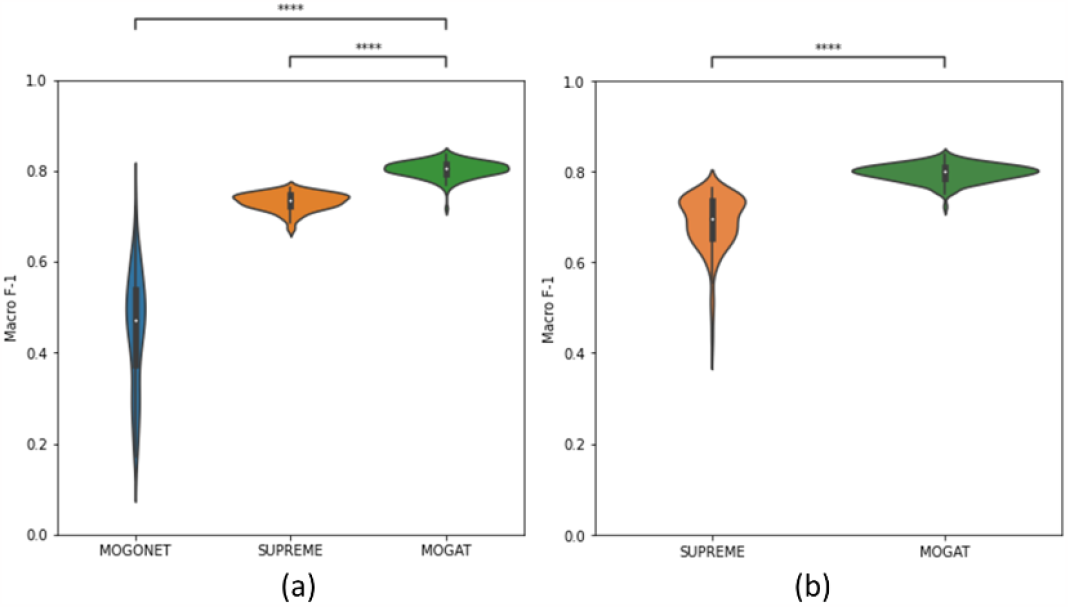
Comparing MOGAT with state-of-the-art multi-omics integration frameworks. Macro-F1 score comparison between (a) MOGONET, SUPREME, and MOGAT using seven combinations of three omics data (EXP, MET, MIR). (b) SUPREME and MOGAT using 255 different combinations of eight omics data. Pairwise statistical comparison was done using Mann-Whitney Wilcoxon test with two-sided Bonferroni correction. The P-values annotations are - ns: 0.05 < p <= 1; *: 0.01 < p <= 0.05; **: 0.001 < p <= 0.01 ***: 0.0001< p <= 0.001 ****: p <= 0.0001

However, for all the omics data, comparison between SUPREME and MOGAT was performed as we found MOGONET incompatible with eight datatypes. The macro-F1 scores on 255 (2^8^-1) different combinations of eight datatypes are calculated and plotted using violin plots in Figure 2(b). It is clear that multi-GAT outperforms SUPREME. We also investigated the contribution of each omics data type in generating GAT embeddings, and results are shown in Table 3. Eight combinations were considered for eight different datatypes, where each combination constitutes all datatypes except the one whose contribution will be investigated. Last row shows the performance of MOGAT using all datatypes. The performance was estimated in terms of accuracy, weighted-F1 score, and macro-F1score. The performance metrics were calculated from ten runs, and their mean and standard deviations are reported. We observed that MOGAT performs better using all types of data than the other eight combinations where one type of data is absent, which means that each data type contributes towards subtype prediction. It is noticeable that the performance without data type EXP (i.e., mRNA expression) is the lowest compared to the performance using all data types with Accuracy 0.837 vs. 0.861, Weighted-F1 score 0.831 vs. 0.861, and Macro-F1 score 0.766 vs. 0.826. This means that mRNA expression contributes the most towards sub-type prediction. On the other hand, the data type MIR (i.e., miRNA expression) has the lowest contribution towards subtype prediction.

**Table 3.**
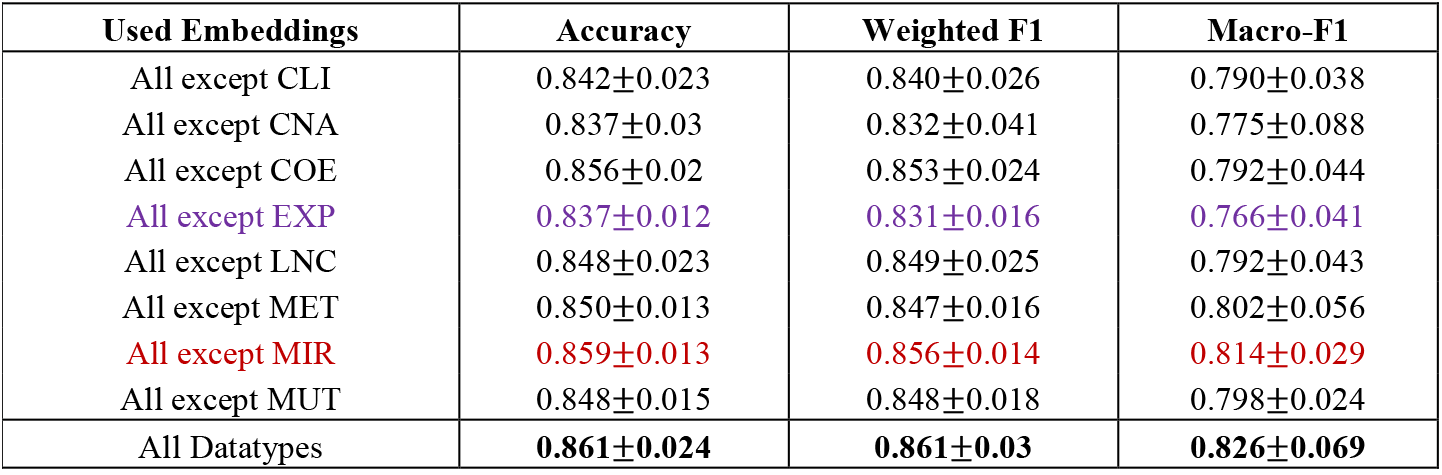
Contribution of individual omics datatype. Performance of MOGAT with different combinations of datatypes in terms of accuracy, weighted-F1, and macro-F1. The last two row contains the result from integrating/embedding all datatypes (eight types). Each of the other rows contains embeddings from every datatype except the one noted in the first column.

| Used Embeddings | Accuracy | Weighted F1 | Macro-F1 |
| --- | --- | --- | --- |
| All except CLI | $0.842 \pm 0.023$ | $0.840 \pm 0.026$ | $0.790 \pm 0.038$ |
| All except CNA | $0.837 \pm 0.03$ | $0.832 \pm 0.041$ | $0.775 \pm 0.088$ |
| All except COE | $0.856 \pm 0.02$ | $0.853 \pm 0.024$ | $0.792 \pm 0.044$ |
| All except EXP | $0.837 \pm 0.012$ | $0.831 \pm 0.016$ | $0.766 \pm 0.041$ |
| All except LNC | $0.848 \pm 0.023$ | $0.849 \pm 0.025$ | $0.792 \pm 0.043$ |
| All except MET | $0.850 \pm 0.013$ | $0.847 \pm 0.016$ | $0.802 \pm 0.056$ |
| All except MIR | $0.859 \pm 0.013$ | $0.856 \pm 0.014$ | $0.814 \pm 0.029$ |
| All except MUT | $0.848 \pm 0.015$ | $0.848 \pm 0.018$ | $0.798 \pm 0.024$ |
| All Datatypes | **$0.861 \pm 0.024$** | **$0.861 \pm 0.03$** | **$0.826 \pm 0.069$** |

### 3.2 Visualizations

To investigate whether the embeddings can capture the underlying insights of the data, we used principal component analysis (PCA) [27] and tSNE [28], which are dimensionality reduction techniques commonly used to reduce the high-dimensional data to a lower-dimensional space and visualize and interpret. Figure 3 shows the PCA and tSNE plot for the learned GAT embeddings with their counterpart raw feature matrix. We observed that the embeddings learned the underlying structure of the data, and the subtype clusters are relatively more distinct than their raw feature counterparts. Clusters of points that denote a breast cancer subtype are more distinguishable in the PCA and tSNE plot of GAT embeddings than their corresponding plot of the raw feature matrix, which was used to train the GATs.

**Fig. 3.**
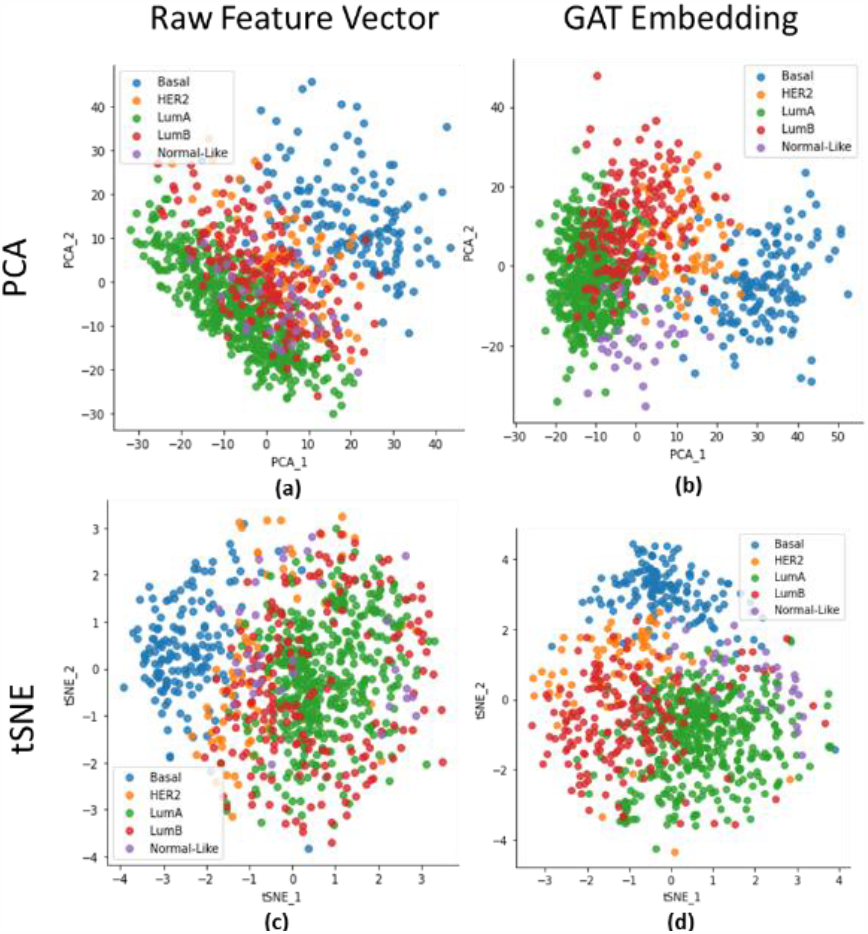
PCA and tSNE plot of breast cancer patients using raw features and embeddings. (a) PCA plot using raw features, (b) PCA plot using GAT embeddings, (c) tSNE plot using raw features, and (d) tSNE plot using GAT embeddings. X-axis and Y-axis correspond to the first and second component of PCA or tSNE.

### 3.3 Survival Analysis

Survival analysis was performed using raw features and GAT embeddings separately, following the methods described in Section 2.3, to evaluate the performance of our framework. The high-risk group contains patients with a risk score higher than the median, and the low-risk group with a score less than or equal to the median. The Kaplan-Meier curves using raw features and GAT embeddings are shown in Figure 4. It is observed that, in both cases, the difference in survival between low-risk and high-risk groups is significant. However, GAT embeddings can significantly distinguish the low-risk and high-risk better than the raw features, as denoted by the log-rank P-value (2.10e^-30^ with GAT embeddings vs. 7.85e^-03^ with Raw features) and hazard ratio (HR: 0.10 with GAT embeddings vs. 0.62 with Raw features).

**Fig. 4.**
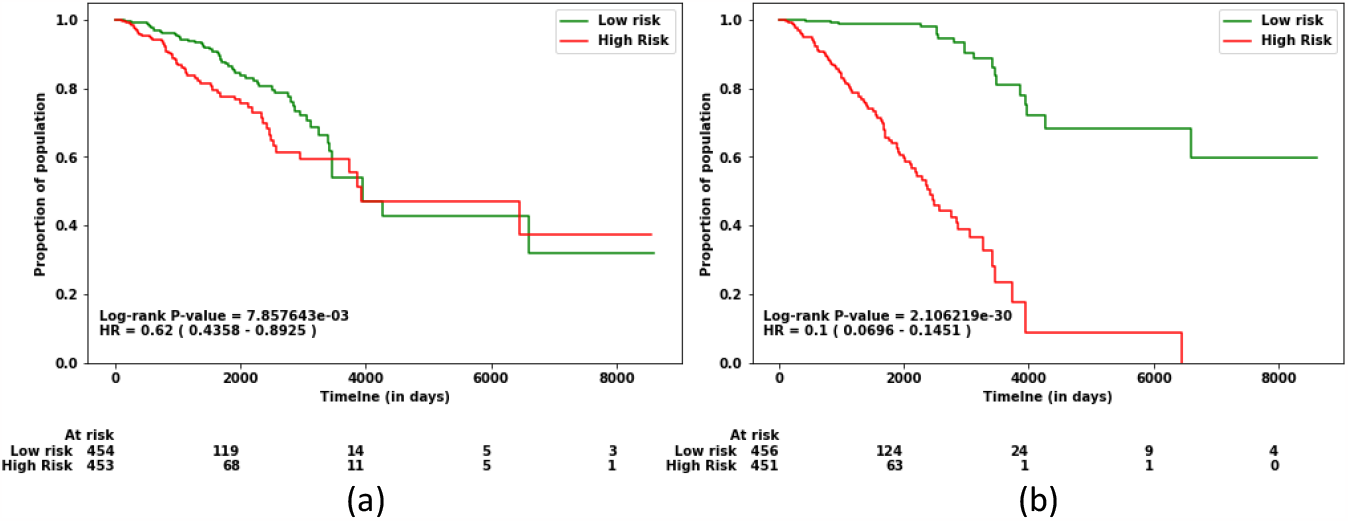
Survival analysis using Raw features and GAT embeddings. Kaplan Meier curve showing the proportion of population of low-risk and high-risk at different observation time. The low-risk and high-risk groups were determined by their risk scores calculated using (a) Raw features and (b) GAT embeddings.

## 4 Discussion

The hypothesis of the present study was that the attention-based graph neural network would provide better embeddings compared to graph convolutional neural network (GCN). Our proposed model, MOGAT, to integrate multi-omics data based on graph attention network (GAT) provides better embeddings and performs better than the GCN based approaches, such as, MOGONET and SUPREME.

The study presented in this work has some limitations that need to be addressed in future works. It was restricted to using the TCGA-BRCA cohort to predict its five different subtypes. The efficacy of the MOGAT framework in predicting subtypes needs to be tested in other cancer types from the TCGA cohort, as well as other diseases, such as, Alzheimer’s and Parkinson’s.

The current study also presents opportunities for future research. The framework used in this study utilized a patient similarity network as the input graph, with each node representing a patient. However, it is possible to reorganize the framework so that each node represents a gene, making the task of the graph neural network a graph classification instead of node classification. While some existing methods follow this approach [5][8], it limits the number of omics data that can be incorporated as node features. For instance, when using genes as nodes, gene expression, somatic mutation, copy number variation, and DNA methylation can be incorporated, but not lncRNAs, miRNAs, and coexpression eigengenes due to the absence of a one-to-one association with genes. To address this, separate graph attention networks with different network and node features are required to incorporate these additional omics data.

## Acknowledge

This work has been supported by the NSF CAREER Award (#1901628) to AMM.

